# Phenotype-specific enrichment of Mendelian disorder genes near GWAS regions across 62 complex traits

**DOI:** 10.1101/324558

**Authors:** Malika Kumar Freund, Kathryn Burch, Huwenbo Shi, Nicholas Mancuso, Gleb Kichaev, Kristina M. Garske, David Z. Pan, Päivi Pajukanta, Gleb Pasaniuc, Valerie A. Arboleda

**Author notes:** Correspondence (MKF). (VAA). These authors jointly supervised this work (Gleb Pasaniuc, Valerie A. Arboleda).

## Abstract

Although recent studies provide evidence for a common genetic basis between complex traits and Mendelian disorders, a thorough quantification of their overlap in a phenotype-specific manner remains elusive. Here, we quantify the overlap of genes identified through large-scale genome-wide association studies (GWAS) for 62 complex traits and diseases with genes known to cause 20 broad categories of Mendelian disorders. We identify a significant enrichment of phenotypically-matched Mendelian disorder genes in GWAS gene sets. Further, we observe elevated GWAS effect sizes near phenotypically-matched Mendelian disorder genes. Finally, we report examples of GWAS variants localized at the transcription start site or physically interacting with the promoters of phenotypically-matched Mendelian disorder genes. Our results are consistent with the hypothesis that genes that are disrupted in Mendelian disorders are dysregulated by noncoding variants in complex traits, and demonstrate how leveraging findings from related Mendelian disorders and functional genomic datasets can prioritize genes that are putatively dysregulated by local and distal non-coding GWAS variants.

## INTRODUCTION

Genetic architectures of human traits have traditionally been classified into two major categories. Typically, complex traits demonstrate polygenic architectures arising from many low-effect common variants, whereas rare traits tend to have high-effect monogenic determinants^1^. The underlying and practical distinction between these classes has historically been based on the presence of highly penetrant, rare, single-gene disruptive mutations causing recognizable clinical monogenic diseases (e.g., cystic fibrosis^2^), and the relative absence of such mutations in complex diseases such as diabetes and schizophrenia^3^. Evidence is accumulating that these two classes of phenotypes may not be as biologically distinct as previously thought^4^. Multiple exceptions to the “common disease, common variant” hypothesis^1^ have been identified for complex traits^5–8^ and their molecular phenotypes^9–12^, and Mendelian disorders have also been found to be affected by multiple or common genetic variants ^13–16^. This suggests that there exists a spectrum of genetic architectures rather than a dichotomous classification. Accordingly, the monogenic forms of complex traits (i.e., phenotypically-matched Mendelian disorders) are increasingly used as a starting point to identify genes relevant to complex traits for further study^17–19^. Furthermore, overlap has been identified between genes linked with Mendelian disorders and genetic determinants of complex traits and diseases such as Parkinson’sdisease^20; 21^, obesity^22^, height^23^, ototoxicity^24^, and others^25^. However, the overlap of each of these complex traits with Mendelian disorders has been examined individually, with different metrics of overlap. In a large study of patient medical records, Blair et al. identified systematic, significant comorbidities between Mendelian disorders and complex diseases, and that association signals from genome-wide association studies (GWAS) for complex diseases were enriched in genomic regions with known roles in comorbid Mendelian disorders, suggesting a shared genetic basis^26^. However, the study focuses on Mendelian disorders comorbid with complex diseases in the same individual, rather than Mendelian disorders demonstrating similar phenotypes to complex traits. Furthermore, advances in sequencing technology have greatly expanded the phenotypic spectrum in known Mendelian syndromes, allowing for deconstruction of syndromic diseases into component medical phenotypes. As such, it is now possible to identify all the component-phenotype consequences of Mendelian disorder genes, allowing for greater resolution in identifying gene-phenotype relationships. However, to the best of our knowledge, no study has taken advantage of this to identify genes causing any related component-phenotype regardless of the Mendelian disorder’s best-known or primary phenotype. Thus, a thorough quantification of the overlap between genes associated with complex traits and genes linked to Mendelian disorders in a phenotype-specific manner remains elusive.

Given that the majority of genome-wide association studies for complex traits and diseases have identified significant associations in non-coding genomic regions^27^, we hypothesize that genes individually involved in Mendelian disease belong to the biological pathway(s) shared by both complex and Mendelian disease. Specifically, we hypothesize that large-effect coding variants disrupt individual genes, resulting in severe phenotypes (i.e., Mendelian disorders), while non-coding variants produce complex traits by collectively dysregulating expression of these same genes, allowing for nuanced or tissue-specific phenotypes. Based on this hypothesis, we expect to identify an enrichment of GWAS signal for a given complex trait near genes linked to Mendelian disorders demonstrating similar phenotypes, but no enrichment near genes linked to Mendelian disorders with phenotypes unrelated to the complex trait of interest. To test this hypothesis, we define “Mendelian disorder genes” as any genes linked to Mendelian disorders in the Online Mendelian Inheritance in Man (OMIM) database^28^, and use the well-curated phenotypic breakdown of Mendelian disorders to identify subsets of these genes linked to particular phenotypes (e.g., growth defects or immune dysregulation) expressed as part of any Mendelian disorder. We then examined publicly available GWAS across 62 complex traits (detailed in Table 1) to identify risk genes (here called “GWAS gene sets”) for each complex trait, and quantified the overlap of each GWAS gene set with 20 other sets of Mendelian disorder genes for particular phenotypes (detailed in Table 1). We find a consistent, significant, and specific enrichment between GWAS gene sets for complex traits and Mendelian disorder genes for matched and related phenotypes (51/1,240 pairs; e.g., rheumatoid arthritis and immune dysregulation), supporting our hypothesis of a shared genetic basis between complex and Mendelian forms of disease. In addition, we observe instances of enrichments between GWAS gene sets for certain complex traits and Mendelian disorder genes for unrelated phenotypes (20/1,240 pairs; e.g., systemic lupus erythematosus and mature-onset diabetes of the young), suggestive of shared biological mechanisms yet to be examined. Furthermore, we find an increase in average effect size of GWAS variants near Mendelian disorder genes for matched phenotypes, and identify examples of associated SNPs found directly at the transcription start sites (TSSs) of these phenotypically-matched Mendelian disorder genes as candidates for functional follow-up. Finally, we report novel examples of significant body mass index (BMI)-associated variants directly interacting with phenotypically-related Mendelian disorder genes *CREBBP* and *CYP19A1,* using human primary white adipocyte-specific Hi-C data^29^. Leveraging the growing body of well-curated phenotypic data from studies of Mendelian disorders, we provide a phenotype-driven approach to identifying genetic pathways shared by Mendelian diseases and complex traits.

**Table 1:**
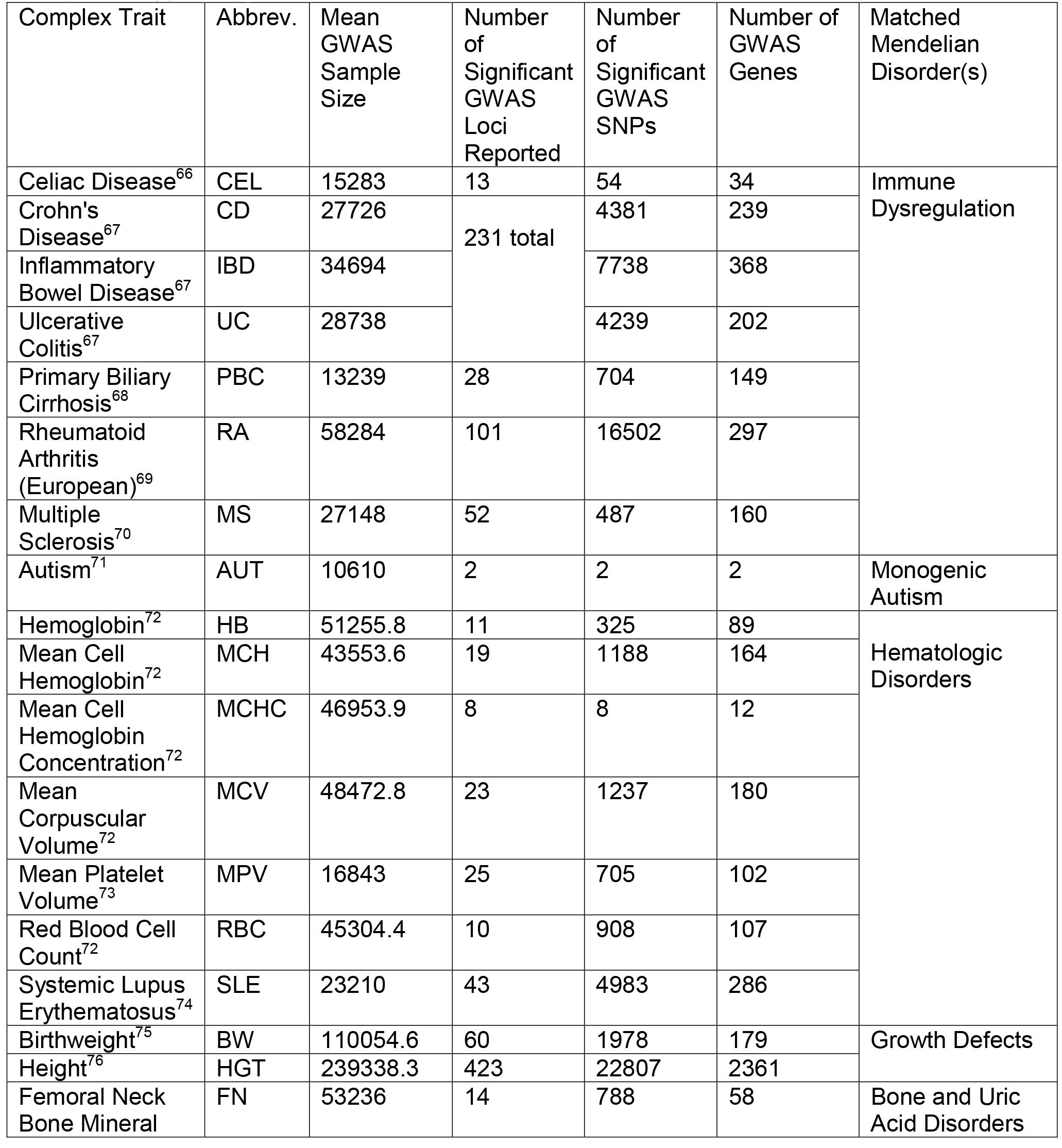
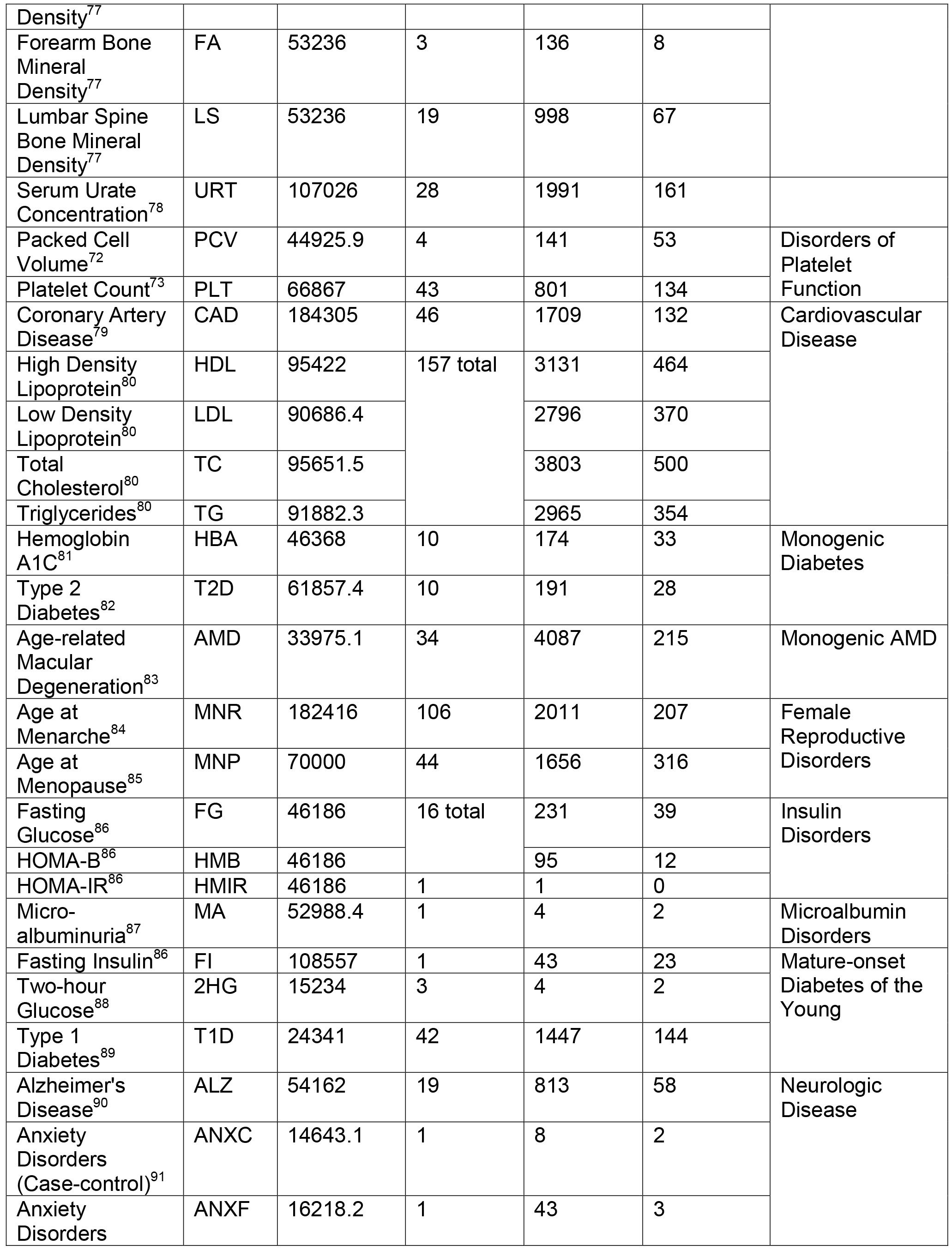
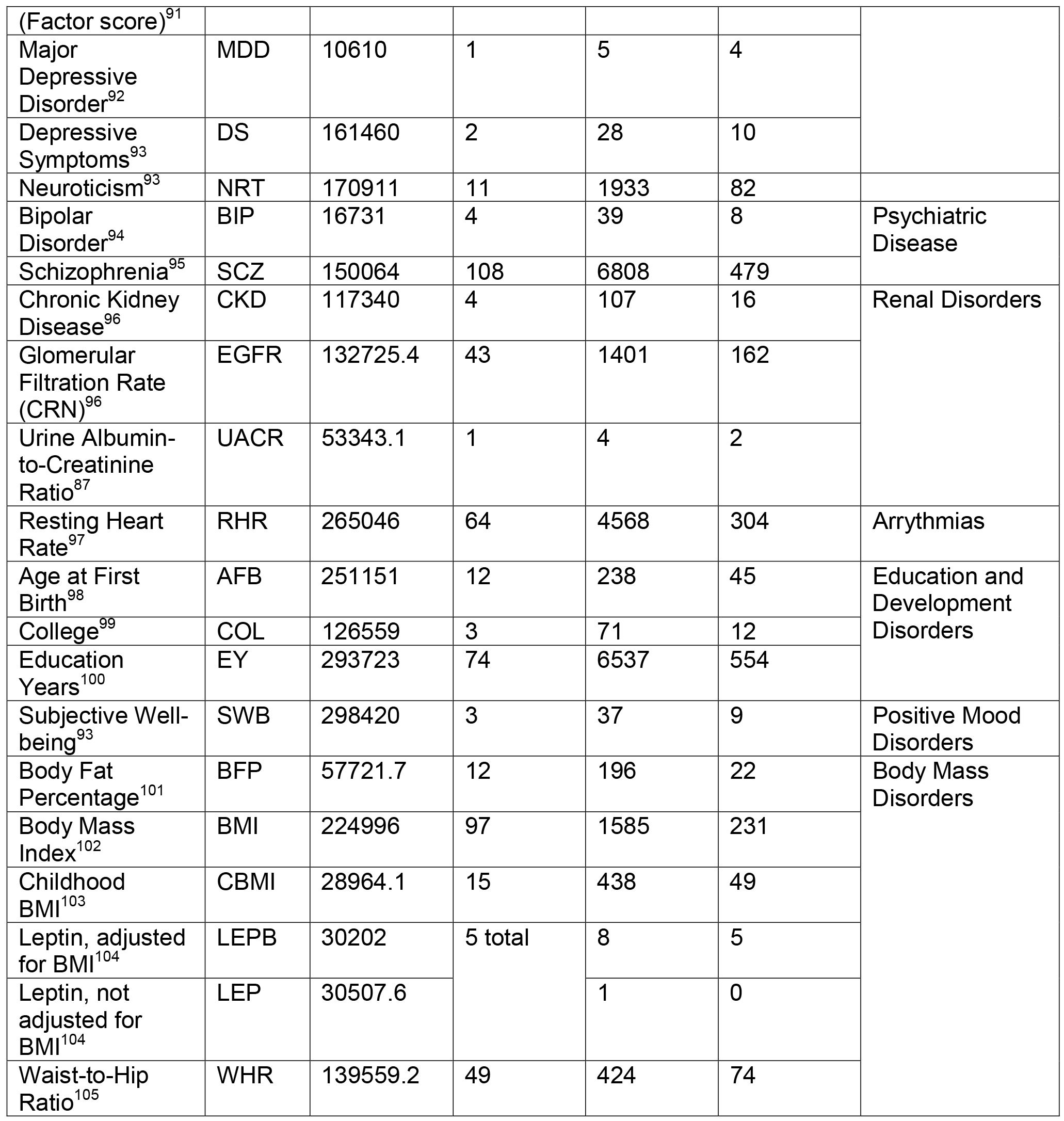
Complex Traits and corresponding Mendelian disorders. This table details the phenotypically-matched pairs of complex traits (N=62) and groups of Mendelian disorders (N=20) examined in our study. Mean GWAS Sample Size and Number of Significant GWAS Loci are reported from original GWAS publications for each complex trait. Significant GWAS SNPs are all SNPs from the publicly available summary statistics meeting genome-wide significance at a threshold of *p* < 5×10^−8^. GWAS genes for each complex trait were identified using the mapping approach described in Methods.

## MATERIAL AND METHODS

### Gene coordinates and symbols

We downloaded gene body coordinates (NCBI build 37/hg19, UCSC Genes track) from the UCSC Table Browser^30^ (see Web Resources) using the gene symbol from the *knownGene* table, transcription start and end sites for each gene from the *knownCanonical* table, and the longest transcript from the *knownGene* table for genes where no entry or multiple entries were listed in the *knownCanonical* table. We used these coordinates for all analyses in our study. Since many genes have been renamed over time, we standardized gene symbols across all analyses in our study by downloading a table of approved symbols, previous symbols, and locus group for each gene from HUGO Gene Nomenclature Committee at the European Bioinformatics Institute (HGNC)^31^ (see Web Resources) and renaming any genes identified by previous symbols with approved gene symbols. We restricted all analyses in our study to genes classified as protein-coding according to the HGNC locus group, from chromosomes 1-22. These processing steps resulted in a final single set of coordinates for 17,695 autosomal protein-coding genes (for data access, see Web Resources).

### Mendelian disorder genes and loss-of-function (LOF) intolerant genes

To identify Mendelian disorder genes, we downloaded the Online Mendelian Inheritance in Man (OMIM) catalogue database^28^ and identified all genes linked to Mendelian disorders satisfying the following criteria: (1) disorder is Mendelian and fully penetrant, therefore excluding susceptibility phenotypes and (2) molecular basis of the Mendelian disorder is known (i.e., phenotype mapping key = 3). We defined loss-of-function (LOF) intolerant genes as any gene with greater than 90% probability of being loss-of-function intolerant, according to the pLI score (pLI > 0.9) from the Exome Aggregation Consortium (ExAC)^32^; this score is derived from the number of observed versus expected LOF variants in a given gene across approximately 60,000 healthy exomes. Following the same restriction and gene symbol standardization criteria described above resulted in a final set of 3,446 Mendelian disorder genes and 2,978 LOF-intolerant genes.

### Phenotype-specific Mendelian disorder gene sets

To identify subsets of Mendelian disorder genes linked to particular phenotypes, for each complex trait we curated a set of standardized clinical phenotype terms to describe the full range of relevant Mendelian phenotypes. We used these terms to search the OMIM database via API for all Mendelian disorders demonstrating these phenotypes, then extracted the gene(s) linked to each Mendelian disorder. We restricted gene-phenotype associations to those satisfying the same criteria (1) and (2) as described above, and with the following additional criteria: (3) gene-phenotype association description does not contain “genome-wide association study” or other GWAS synonyms unless: the description also contains any of the terms “missense”, “nonsense”, “nonsynonymous”, or “frameshift”; or the gene contains at least one pathogenic or likely pathogenic allele in the ClinVar database^33^. We include a full list of phenotype-specific Mendelian disorder gene sets and clinical phenotype terms used in **Table S1**.

A comparison of all phenotype-specific Mendelian disorder gene sets revealed a high degree of overlap among the gene sets for clinically-related Mendelian phenotypes (**Figure S1**). Accordingly, we clustered gene sets based on pairwise overlap, and intersected gene sets visually clustering together to create a single gene set for the representative group of Mendelian disorders. After combining similar gene sets, a total of 20 non-disjoint phenotype-specific Mendelian disorder gene sets remained with an average of 375 genes per set; we include a description of each cluster in **Table S2**.

### Complex trait gene sets

We downloaded publicly available summary statistics (per-allele SNP effect sizes, or log-odds ratios for case-control traits, with standard errors^34^) for large-scale GWAS of 62 traits^26^ (Table 1; average N=83,170; some GWAS were imputed using the 1000 Genomes Project as a reference panel by their respective consortia while others were not.) For each trait, we identified a gene set by mapping each autosomal genome-wide significant SNP (*p* < 5×10^−8^) to the closest up- and downstream protein-coding genes as defined above, resulting in a total of 62 non-disjoint GWAS gene sets. As GWAS regions often contain multiple genome-wide significant SNPs, and the true causal gene may not lie adjacent to the lead SNP in a region^35; 36^, we defined GWAS gene sets by mapping genes with respect to every genome-wide significant SNP rather than only the index GWAS SNPs at each genomic risk region.

### Quantifying overlap between complex trait and Mendelian disorder

For each complex trait-Mendelian disorder pair, we compared the GWAS gene set and phenotype-specific Mendelian disorder gene set using a 2×2 contingency table (counting whether each gene was in the GWAS gene set or not, and in the Mendelian disorder gene set or not), with the set of autosomal protein-coding genes (n=17,695) representing the total sample. We used Fisher’s Exact Test^37^ to determine significance. Phenotype-specificity of overlap significance was assessed by comparing the GWAS gene sets for each complex trait (n=62) to all phenotype-specific Mendelian disorder gene sets (n=20), a total of 1,240 pairs. Significance was assessed at a trait-specific Bonferroni-corrected threshold (*p* < 0.05/20) based on the number of phenotype-specific Mendelian disorder gene sets (n=20) being compared to each GWAS gene set. Since not all autosomal protein-coding genes are Mendelian, we also assessed significance in permutations; we drew 10,000 random sets of Mendelian disorder genes and 10,000 random sets of any protein-coding genes with replacement (matching the size of each random gene set to the size of the phenotype-specific Mendelian disorder gene set), and counted the number of times more genes were shared between the GWAS gene set and a random set.

To assess the robustness and stability of our SNP-gene mapping approach for complex traits, we performed an overlap quantification with phenotype-specific Mendelian disorder genes using GWAS gene sets derived from two additional SNP-gene mapping methods: by mapping each SNP to all genes within a 50Mb window, and to all genes within a 500Mb window. Comparison of the odds ratios produced by Fisher’s Exact Test for the comparisons of GWAS gene sets (derived by each mapping method) and phenotype-specific Mendelian disorder gene sets demonstrates no major difference in outcomes from different mapping methods (**Table S3**).

### Estimating enrichment of GWAS SNP association signal

We created genomic annotations to capture the regions spanning 50kb upstream through 50kb downstream of gene bodies for four categories of genes: all protein-coding genes (N=17,695), all Mendelian disorder genes (N=3,446), all LOF-intolerant genes (N=2,978), and the phenotype-specific Mendelian disorder gene sets (average N=609). For each complex trait-gene category pair, we computed enrichment of GWAS signal within the category *c* with respect to the set of all protein-coding genes as

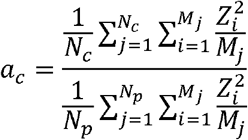

where *N*_*c*_ is the number of genes in category *c, M*_*j*_ is the number of SNPs within 50kb of gene *j, Z*_*i*_ = GWAS effect size of SNP *i* divided by standard error, with total number of protein-coding genes *N*_*p*_. Thus, *a*_*c*_ is the enrichment in average SNP effect size (Z^2^) per gene in category (compared to average Z^2^ for any protein-coding gene). The percent increase in average SNP effect size per gene for category *c*, or (*a*_*c*_ – 1) * 100, is shown in Figure 3. We performed similar comparisons for median SNP effect size per gene for category *c,* and maximum SNP effect size per gene for category *c* (**Table S4**).

To ensure that this signal was not driven by linkage disequilibrium (LD), minor allele frequency (MAF), or average gene length per category, we compared these three properties across the gene categories for each complex trait. We calculated LD scores^38^ reflecting the amount of LD tagged by each SNP in the HapMap 3 reference panel^39^. For each gene category, we averaged the LD scores of SNPs falling within 50kb of each gene, and found no consistent difference across gene sets. Similar analyses were performed to examine average MAF per gene and average gene length per category across each complex trait (**Table S5**); we again found no consistent difference across gene sets.

### Putative causal mechanisms at GWAS risk regions

We performed statistical fine-mapping of the genome-wide significant regions (*p* < 5 × 10^−8^) for each GWAS using fgwas^40^ with no functional annotations and default parameter settings. For each GWAS, we constructed a 95% credible set (defined as the minimum set of SNPs where 95% of the probability of causation at a region is accumulated) for each 5kb region (as default assigned by fgwas) containing a significant GWAS association. We achieved this by adding SNPs one at a time with a decreasing posterior probability of causation (posterior probability of association for the SNP, conditioned on there being an association in the region) until a cumulative 95% probability of causation is reached.

### Identification of candidate regulatory variants

We intersected credible sets for each complex trait with genomic regions 1kb upstream of each phenotypically-relevant Mendelian disorder gene to identify SNPs localizing at the TSS. To identify candidate regulatory variants interacting with promoters of phenotype-matched Mendelian disorder genes, we used interactions from promoter capture Hi-C in human primary white adipocytes ^29^ for each complex trait, and filtered interactions to pairs of interacting regions where at least one region contained a promoter of a phenotype-specific Mendelian disorder gene. We then intersected interaction pairs for each of these regions with credible sets for each complex trait to identify credible SNPs interacting with regions containing promoters of phenotype-specific Mendelian disorder genes.

## RESULTS

### GWAS risk genes show specific, significant overlap with phenotypically-matched Mendelian disorder genes

We first sought to examine the degree of overlap between phenotype-matched Mendelian disorder genes with risk genes for complex traits as identified through GWAS. For each complex trait, we identified corresponding Mendelian forms, often as familial forms or rare phenotypic extremes, and curated Mendelian disorder gene sets composed of Mendelian disorder genes known to cause those specific phenotypes from the OMIM database^28^ (see Methods, and Figure 1). We combined similar Mendelian disorder gene sets to create one gene set for the representative Mendelian disorder(s) (for a total of 20 Mendelian disorder gene sets). We separately ascertained GWAS gene sets for each complex trait by identifying the closest up- and downstream genes to each GWAS SNP meeting genome-wide significance (see Methods, and Figure 1). Overlap between each phenotype-specific Mendelian disorder gene set (n=20) and each GWAS gene set (n=62) was assessed using Fisher’s Exact Test, for a total of 1,240 comparisons (Table 1 and **Table S6**). We hypothesized that GWAS gene sets would have a specific significant enrichment of Mendelian disorder genes for matched Mendelian disorders, but no enrichment for unrelated Mendelian disorders. Among all 1,240 pairs of complex and phenotype-specific Mendelian disorder gene sets assessed, we identified 71 pairs with significant overlap at a *p* < 0.05/20 (Figure 2). An examination of the log-odds ratios for each overlap comparison revealed more extreme enrichments among phenotypically-matched pairs compared to phenotypically-unmatched pairs (Table 2), which is consistent with our hypothesis. 51 out of the 71 significantly overlapping pairs showed perfectly matching phenotypes or reflected known shared biology. Specifically, in many of these pairs, monogenic forms of the complex trait have been well established in the genetics literature; examples include Age-related Macular Degeneration (AMD) and cholesterol traits (high-density lipoprotein (HDL), low-density lipoprotein (LDL), total cholesterol (TC), and triglycerides (TG))^6; 41–44^. We confirmed significant enrichment after trait-specific Bonferroni adjustment (20 comparisons, see Methods) between many of these previously reported pairs such as the complex and monogenic forms of height^45^ (OR=1.39, p=0.029) and HDL and Mendelian forms of cardiovascular disease^46^ (OR=2.10, p=0.006). We also identified previously unreported enrichments; for example, we find a strong enrichment between inflammatory bowel disease (IBD) and Mendelian forms of immune dysregulation (OR=3.32, p=0.01) and between hemoglobin (HB) and Mendelian hematologic disorders (OR=3.99, p=0.009). The remaining 20 pairs with significant overlap suggested novel shared biological mechanisms between complex traits and Mendelian disorders yet to be established (Table 3). For example, we observed an enrichment between height and renal disorders (OR=1.48, p=0.001), and enrichment between Crohn’s Disease and mature-onset diabetes of the young (OR=2.69, p=0.005). Of note, the vast majority of complex and Mendelian disorder gene set pairs without perfectly-matched phenotypes (n=1,158 of 1, 178) did not demonstrate any significant overlap (**Table S6**).

**Figure 1:**
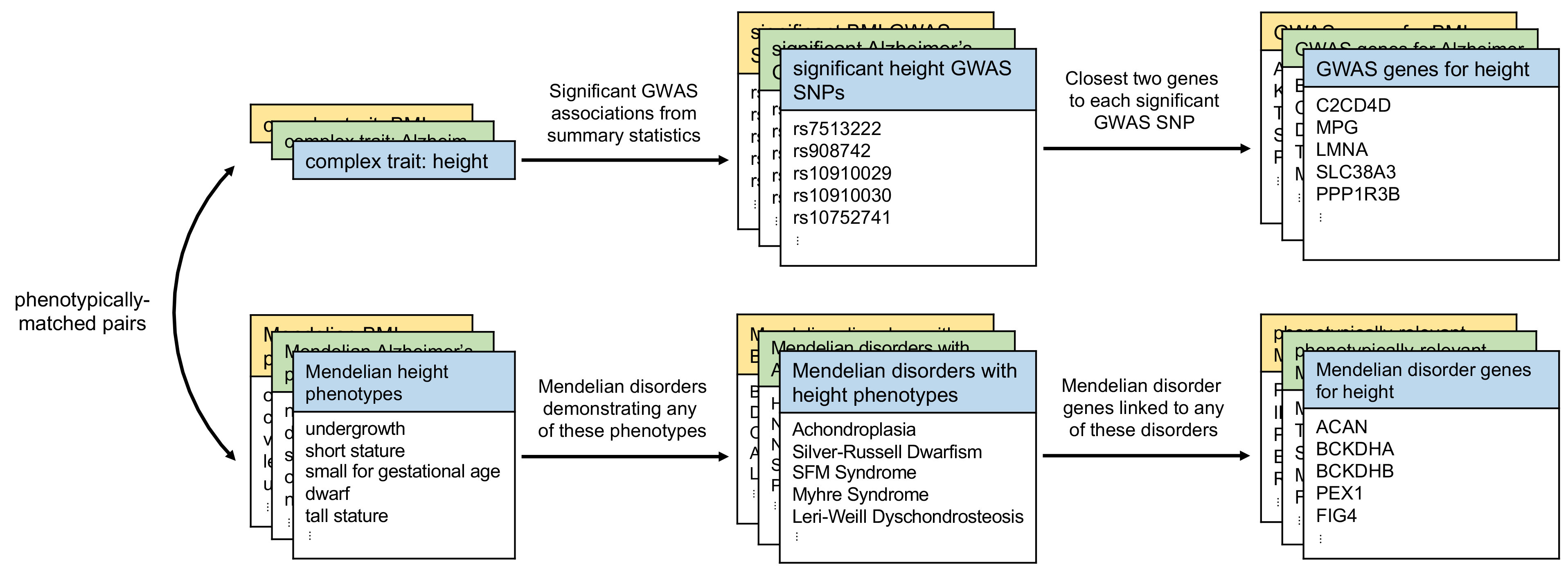
GWAS gene sets and phenotype-specific Mendelian disorder gene sets. For each complex trait (e.g., height), we first identified matched Mendelian phenotypes (e.g., undergrowth, short stature; **Table S9**). Using publicly available GWAS data, we defined the “GWAS genes” for a given complex trait to be the closest upstream and closest downstream protein-coding gene for every genome-wide significant variant in the GWAS. We selected phenotype-matched Mendelian disorder genes by first identifying Mendelian disorders expressing any of the matched Mendelian phenotypes, and then identifying all genes causing any of those disorders.

**Figure 2:**
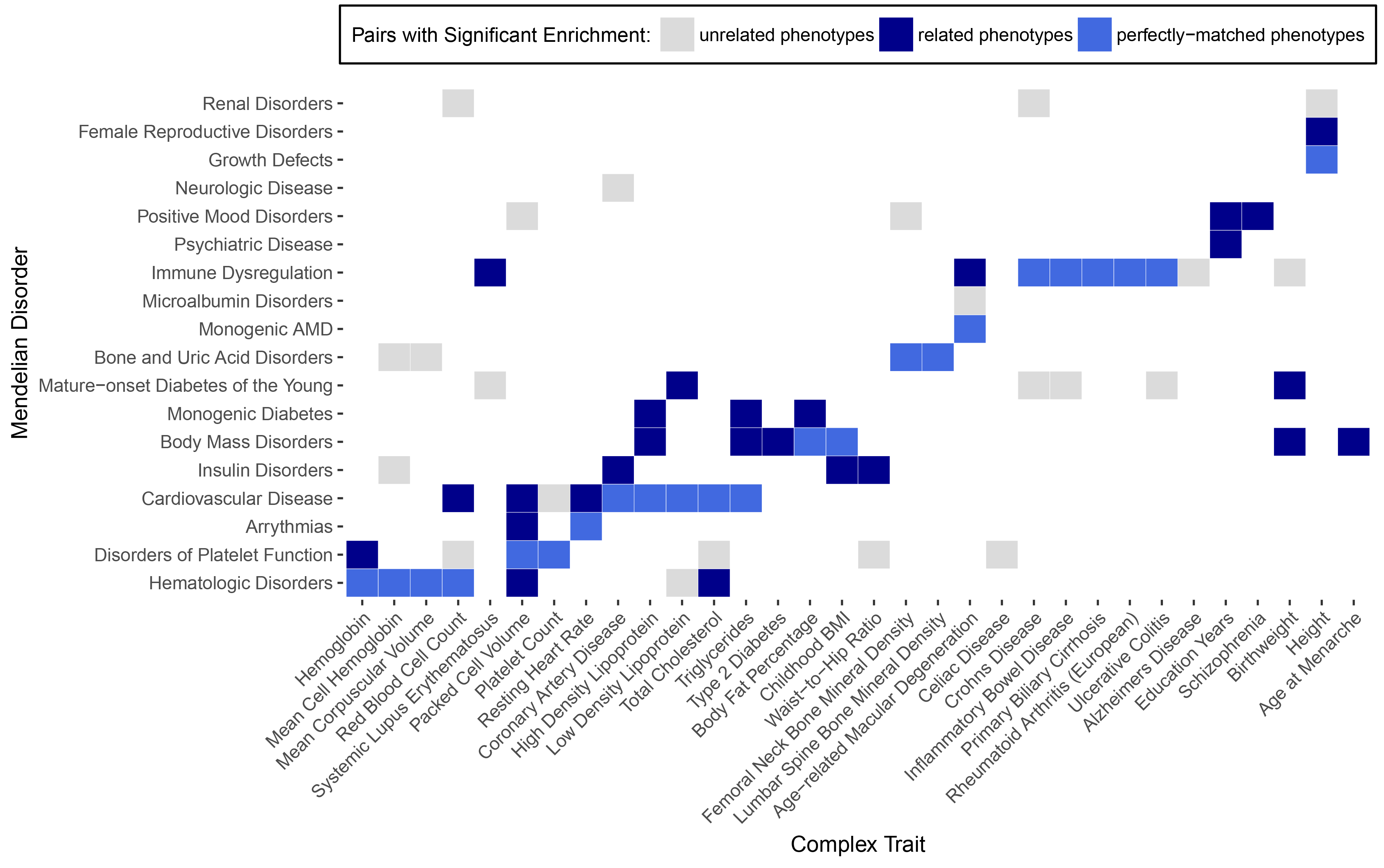
Overlap of GWAS genes with unrelated Mendelian disorder genes demonstrates trait-specificity. Significant overlaps from phenotypically-matched pairs of complex traits and Mendelian disorders (blue) and pairs with unrelated phenotypes (grey) are shown. Phenotypically-matched pairs are subdivided into pairs with perfectly-matched phenotypes (light blue) and pairs with related phenotypes (dark blue). Traits with no significant overlaps are excluded. Significance was assessed at a threshold for each complex trait (*p* < 0.05/20) based on the number of Mendelian disorder gene sets (n=20) compared to each GWAS gene set.

**Table 2:**
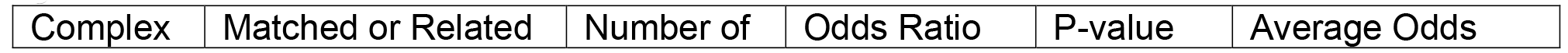
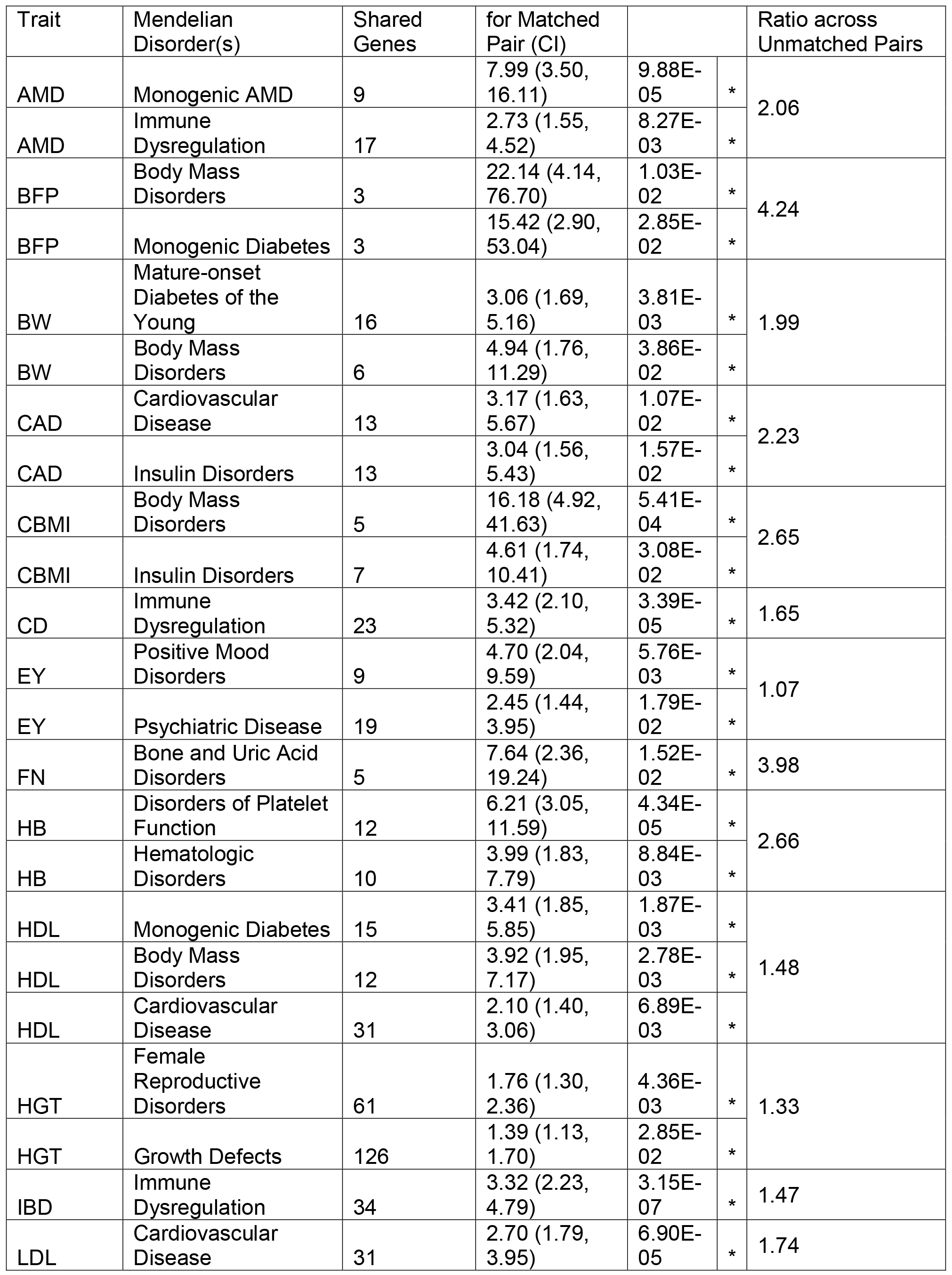
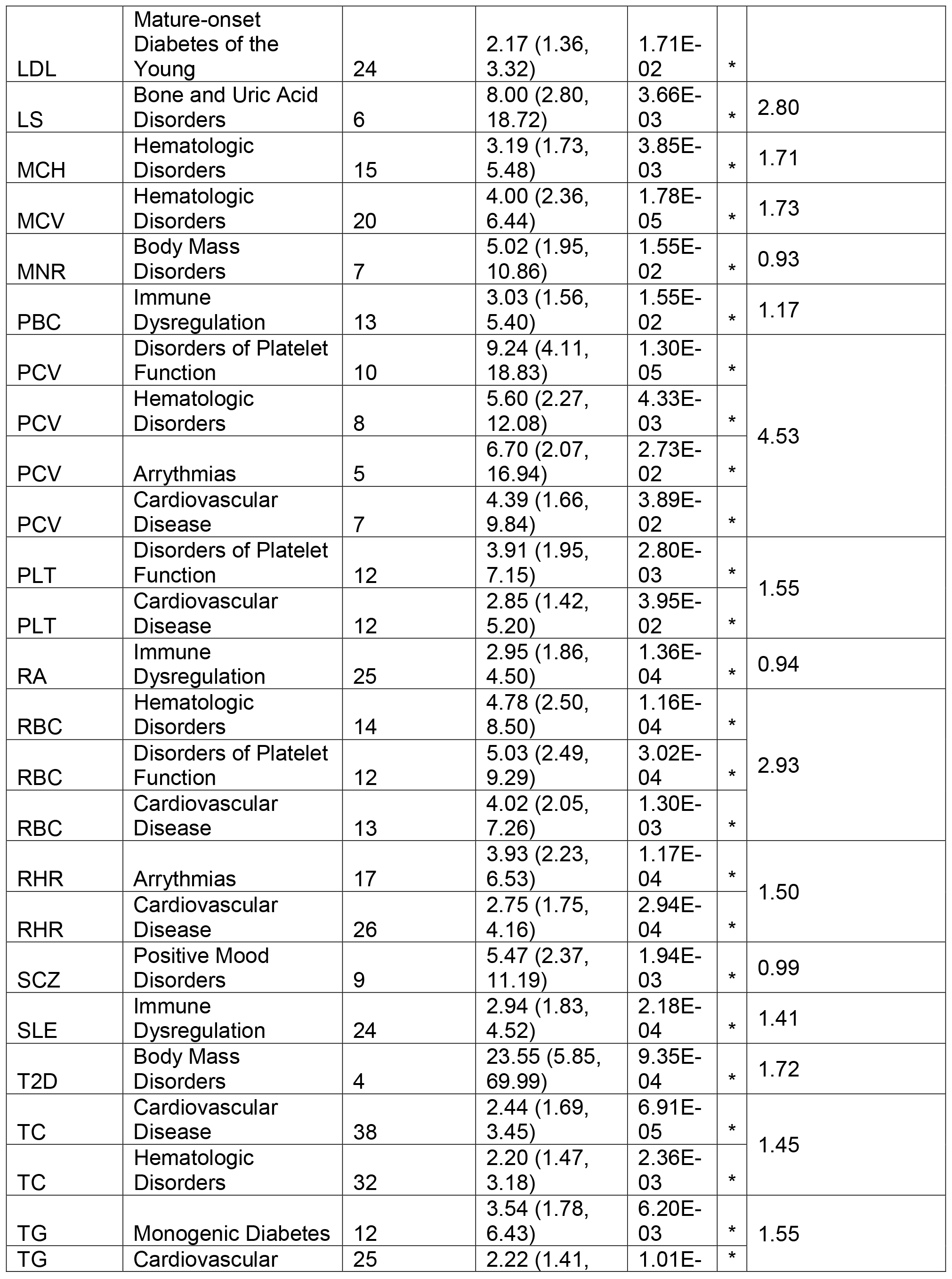
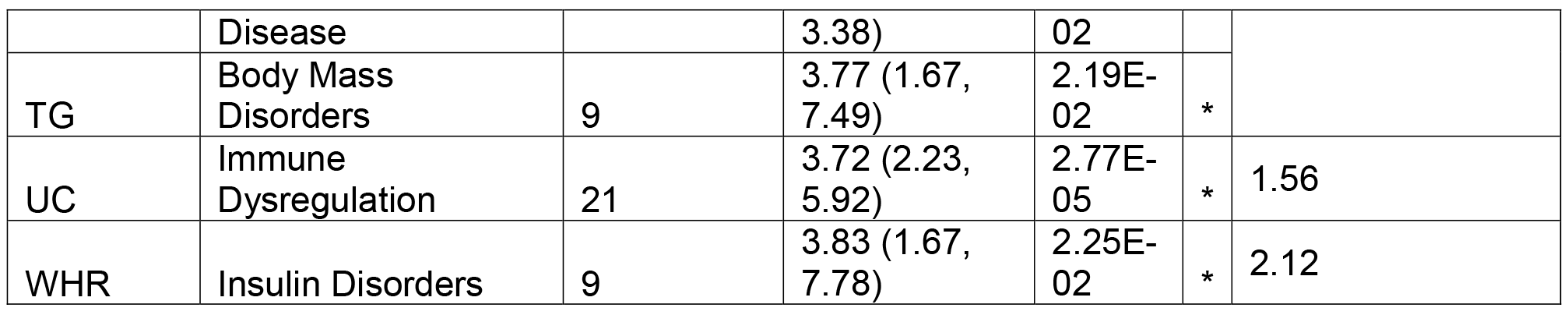
Overlap of GWAS genes and phenotypically-matched Mendelian disorder genes. For each pair of complex trait and Mendelian disorder, Fisher’s exact test was used to quantify the enrichment of shared genes with an odds ratio and p-value (see Methods). Significance was assessed at a threshold of *p* < (0.05/20) correcting for the number of Mendelian disorder gene sets compared to each complex trait gene set. This table lists pairs of complex traits and phenotypically-matched or related Mendelian disorders with significant overlap. For comparison, the average odds ratio for pairings of each complex trait with all unrelated Mendelian disorder gene sets is included.

**Table 3:**
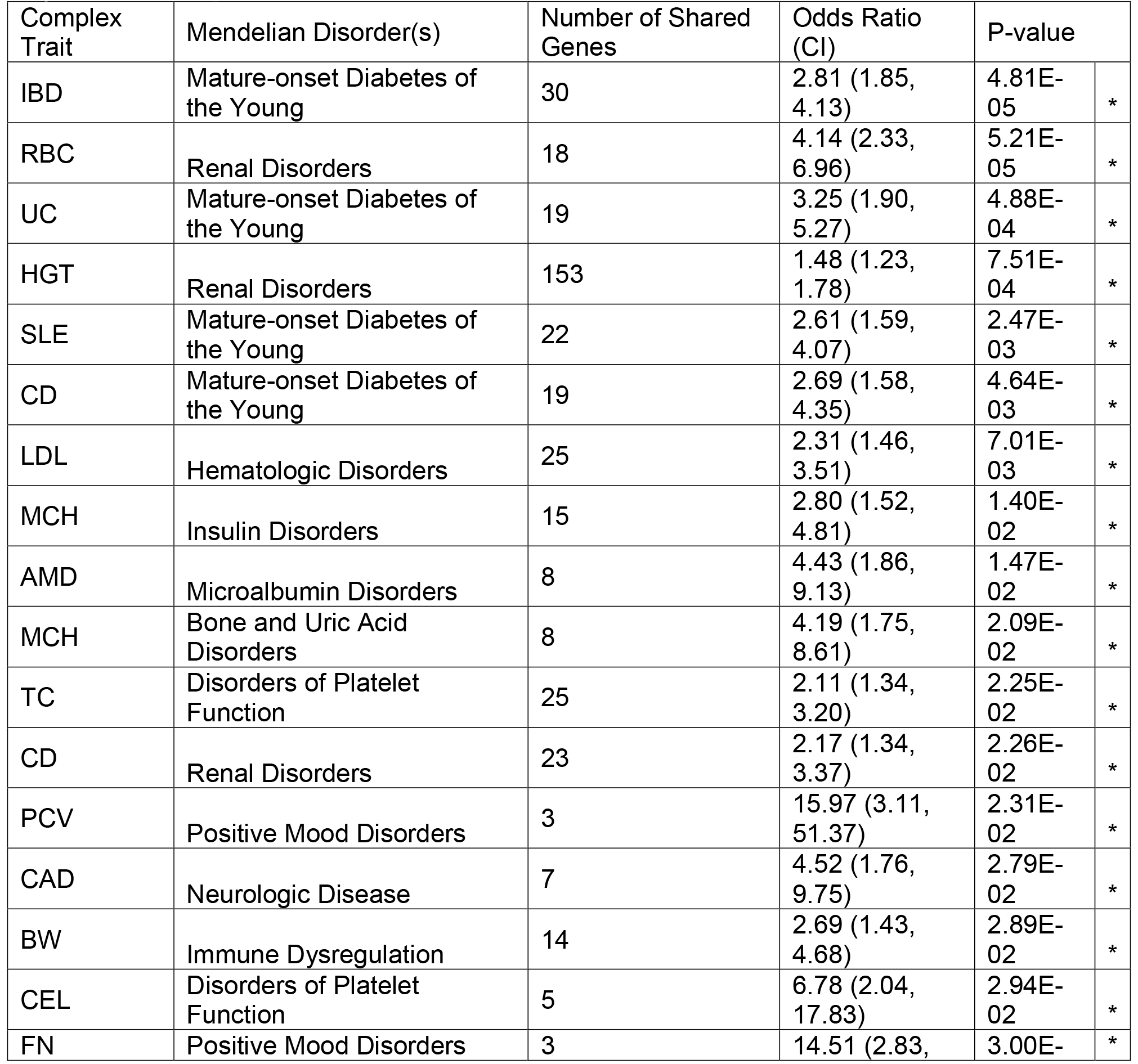
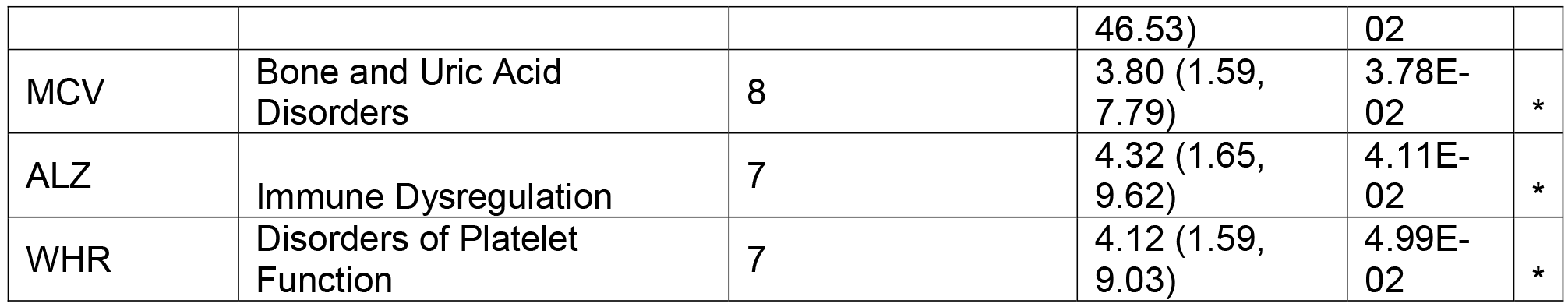
Instances of significant overlap of GWAS genes and unrelated Mendelian disorder genes. As in Table 2, Fisher’s exact test was used to quantify the enrichment of shared genes between complex traits and Mendelian disorders with an odds ratio and p-value (see Methods). Significance was assessed at a threshold of *p* < (0.05/20) correcting for the number of Mendelian disorder gene sets compared to each complex trait gene set. This table lists pairs of complex traits and phenotypically-unrelated Mendelian disorders that demonstrated significant overlap.

### SNPs near phenotypically-matched Mendelian disorder genes show increased effect size on complex traits

Because Mendelian disorder genes exhibit severe biological effects when either one or both alleles are disrupted, dysregulation of the gene through changes in expression or other mechanisms might have a more significant effect than dysregulation of a non-Mendelian disease gene. We hypothesized that SNPs near these phenotype-specific Mendelian disorder genes have further increased effects on complex traits due to the increased biological relevance of these gene categories. From the publicly available GWAS summary statistics for each complex trait, we computed the average GWAS effect sizes of SNPs falling within each protein-coding gene, and compared the average effect sizes per gene across all Mendelian disorder genes and across phenotypically-relevant Mendelian disorder genes (see Methods). Across complex traits, we found an increased average effect size per gene for all Mendelian disorder genes and a further increased average effect size per gene for phenotypically-relevant Mendelian disorder genes (Figure 3 and **Table S4**). This suggests that the genomic regions containing the most biologically-relevant genes for each trait contribute most significantly to complex trait biology. We also confirmed that loss-of-function (LOF) intolerant genes (as defined by ExAC’s pLI score > 0.9, see Methods) demonstrate a higher average effect size across most complex traits examined^32^. Given the extreme intolerance of deleterious mutations in these genes, it is possible that LOF-intolerant genes demonstrate embryonic lethal mutant phenotypes, and are thus undiscovered as Mendelian disorder genes at this time. We found no significant increase in linkage disequilibrium or decrease in average minor allele frequency (MAF) of the SNPs within each category compared to the SNPs within all protein-coding genes (**Table S5**), suggesting that the observed signal is not driven by any of these confounders. Of note, we did observe a respective increase in average gene length between all protein-coding genes, all Mendelian disorder genes, and LOF-intolerant genes (**Table S5**). Therefore, it is possible that our findings of enriched GWAS signal in these gene categories is due instead to longer genes being more likely to tag causal variation. However, in general we did not observe a significant increase in average gene length for the phenotype-specific Mendelian disorder gene sets as compared to all Mendelian disorder genes (**Table S5**), but still found an increase in enrichment of GWAS signal (Figure 3), suggesting that gene length is not significantly confounding our results. The only exceptions to this are the phenotype-specific Mendelian disorder gene sets for neurological phenotypes, for which the average gene length was increased compared to all Mendelian disorder genes; this is consistent with other reported findings about gene length in neurological traits^47^.

**Figure 3:**
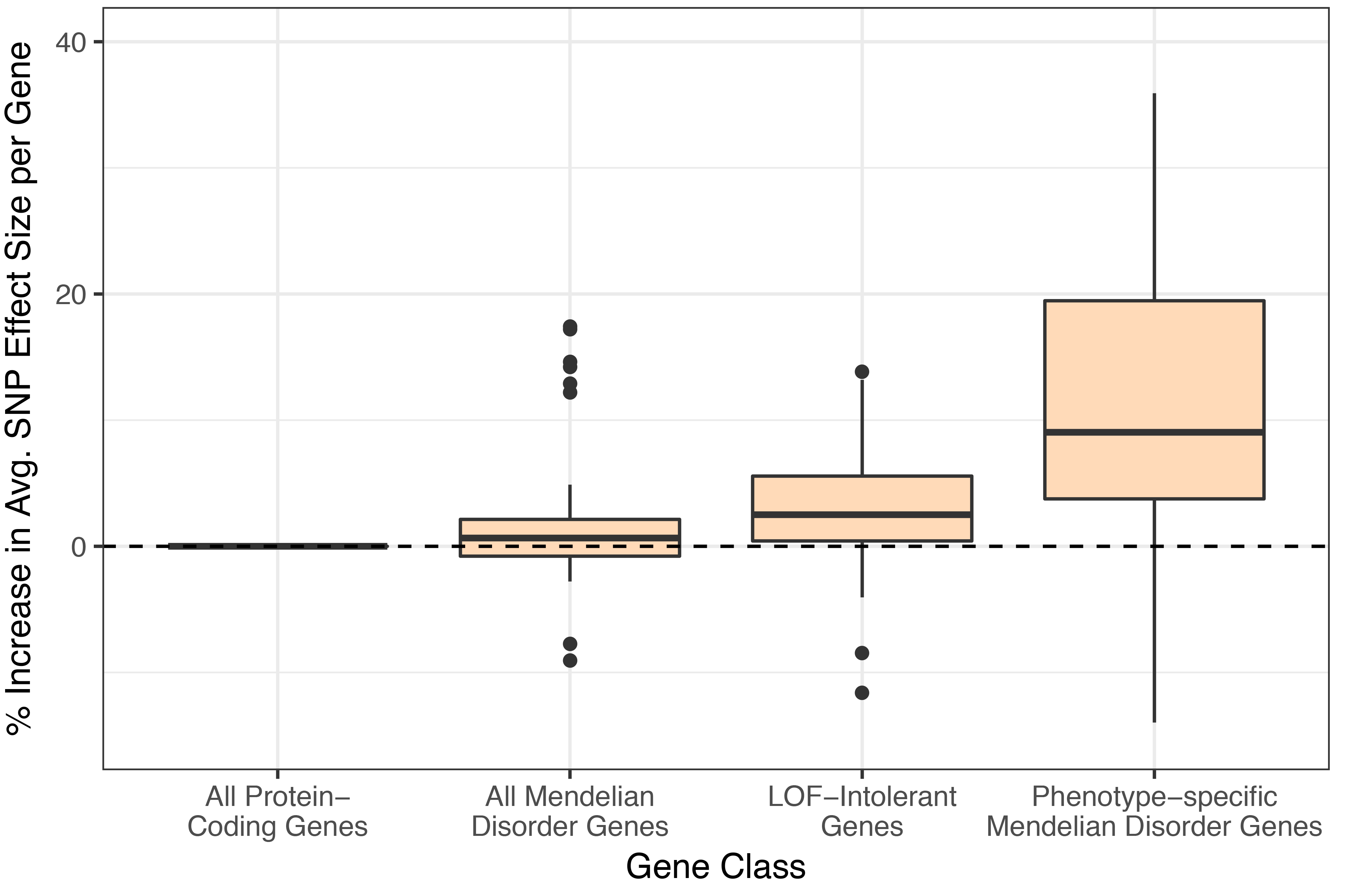
Effect sizes for SNPs on complex traits from GWAS are increased for genes that are loss-of-function intolerant and for phenotypically-relevant Mendelian disorder genes. The increase in average SNP effect size per gene across four gene categories. We averaged effect size (Z^2^) across all SNPs falling within 50kb of a gene to obtain an average SNP effect size per gene, and averaged across all genes in each category (all protein coding genes, all Mendelian disorder genes, all LOF-intolerant genes, and all phenotypically-relevant Mendelian disorder genes for each trait). We normalized these averages to the average SNP effect per gene for any protein coding genes. The box plots represent the distribution of increase in average effect size per gene across all traits.

### Examples of credible SNPs for GWAS regions near phenotypically-matched Mendelian disorder genes

We next sought to identify common non-coding variants that may causally impact complex trait phenotypes by dysregulating phenotypically-relevant Mendelian disorder genes. For each complex trait, we performed statistical fine-mapping of significant GWAS regions to construct 95% credible sets for each region (see Methods), and identified SNPs from the credible set located at the TSS of a gene from the phenotypically-relevant Mendelian disorder gene set. We found a total of 786 credible set SNPs (out of approximately 3.5 million) localizing at the TSS of a phenotypically-relevant Mendelian disorder gene (an average of 20 SNPs per trait, for 38 traits where at least one such SNP was found; **Tables S7 and S8**), and identified 25 promising candidate SNPs (attaining genome-wide significance in GWAS) at TSSs that could be regulating the proximal Mendelian disorder gene (Table 4). We highlight two examples: first, we found a significantly associated SNP from the credible set for coronary artery disease (rs1332327, Z=6.798) at the promoter of *LIPA* (MIM# 278000), a Mendelian disorder gene linked to Wolman Disease and Cholesteryl Ester Storage Disease (both Lysosomal Acid Lipase Deficiencies, MIM# 278000) causing hypercholesterolemia and hypertriglyceridemia as part of cholesteryl ester- and triglyceride-filled macrophage infiltration syndromes (Figure 4A). Second, from the credible set for red blood cell count, we found a significantly associated SNP (rs1010222, Z= −5.961) at the promoter of *CALR* (MIM# 109091), a Mendelian disorder gene known to cause Myelofibrosis (MIM# 254450) involving generalized bone marrow fibrosis, reduced hemopoiesis, no hemophagocytosis, and myeloproliferative disease (Figure 4B). In both cases, the putative causal SNP for the complex trait lies immediately upstream of the TSS of the phenotypically-relevant Mendelian disorder gene, in addition to falling within regions containing by regulatory epigenetic marks.

**Figure 4:**
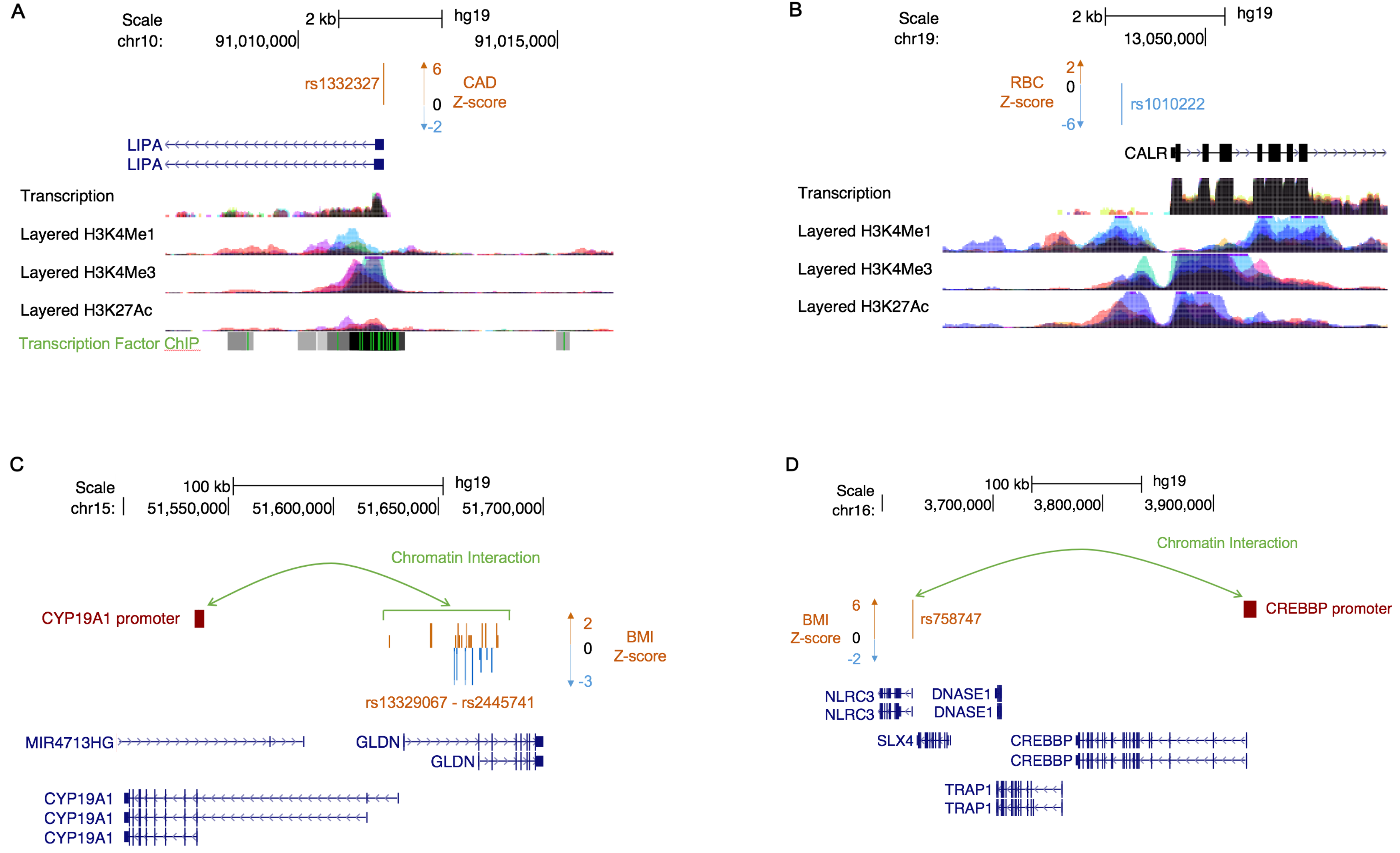
Candidate regulatory SNPs fall at transcription start sites and long-range promoters of phenotypically-relevant Mendelian disorder genes. **A, B)** Shown here are two examples of putative causal SNPs localizing at a TSS of a phenotypically-relevant Mendelian disorder gene. **A**) Putative causal SNP rs1332327, associated with coronary artery disease (Z = −5.961), lies at the TSS of *LIPA.* **B**) Putative causal SNP rs1010222, associated with red blood cell count with a Z-score of −5.961, lies at the TSS of *CALR.* **C, D)** Shown here are two representations of chromatin interactions in white adipose tissue. **C**) A cluster of SNPs from the credible set of variants associated with BMI (Z-score plotted in orange and blue) physically interacts with the promoter of a particular isoform of *CYP19A1*. **D**) A single SNP (rs758747) from the credible set, associated with BMI (Z = 6.081), physically interacts with the promoter of a distant gene *CREBBP.*

**Table 4:** Genome-wide significant SNPs localizing at TSS of phenotypically-relevant Mendelian disorder genes. GWAS SNPs from the credible set for each complex trait were intersected with transcription start site (TSS) regions 1kb upstream of phenotypically-matched Mendelian disorder genes. This table lists all genome-wide significant SNPs (*p* < 5×10^−8^ from GWAS, with chromosomal location) from all complex traits localizing at the tSs of a phenotypically-matched Mendelian disorder gene (italicized).

| Complex Trait | SNP ID | Chr | Position | Z score | Gene | Strand |
| --- | --- | --- | --- | --- | --- | --- |
| PBC | rs13239597 | chr7 | 128695983 | 9.85309 | *TNPO3* | - |
| HGT | rs8028537 | chr15 | 89345947 | -9.333 | *ACAN* | + |
| HGT | rs10853751 | chr19 | 41903220 | 8.71 | *BCKDHA* | + |
| CD | rs59283234 | chr5 | 150225587 | -8.454 | *IRGM* | + |
| CD | rs751627 | chr5 | 150225113 | -8.451 | *IRGM* | + |
| CD | rs35707106 | chr5 | 150225377 | -8.332 | *IRGM* | + |
| HGT | rs2298307 | chr6 | 80816296 | 8.276 | *BCKDHB* | + |
| HGT | rs12386601 | chr7 | 92157886 | 8.2 | *PEX1* | - |
| BMI | rs17066842 | chr18 | 58040624 | -7.542 | *MC4R* | - |
| HGT | rs12192268 | chr6 | 110011458 | -7 | *FIG4* | + |
| CAD | rs1332327 | chr10 | 91011681 | 6.798 | *LIPA* | - |
| RA | rs13239597 | chr7 | 128695983 | 6.65672 | *TNPO3* | - |
| IBD | rs59283234 | chr5 | 150225587 | 6.51 | *IRGM* | + |
| IBD | rs751627 | chr5 | 150225113 | 6.507 | *IRGM* | + |
| HGT | rs7592246 | chr2 | 219926221 | 6.452 | *IHH* | - |
| IBD | rs34005003 | chr5 | 150225199 | 6.427 | *IRGM* | + |
| IBD | rs35707106 | chr5 | 150225377 | 6.326 | *IRGM* | + |
| MNR | rs3775971 | chr4 | 104641920 | 6.20413 | *TACR3* | - |
| IBD | rs27741 | chr16 | 28504181 | 6.109 | *CLN3* | - |
| HGT | rs4244808 | chr11 | 2163110 | 6.061 | *IGF2* | - |
| RBC | rs1010222 | chr19 | 13048608 | -5.96154 | *CALR* | + |
| CD | rs27741 | chr16 | 28504181 | -5.866 | *CLN3* | - |
| HGT | rs613924 | chr11 | 65769295 | -5.862 | *BANF1* | + |
| AFB | rs4845357 | chr1 | 153896212 | -5.775 | *GATAD2B* | - |
| HGT | rs6591226 | chr11 | 66675990 | 5.517 | *PC* | - |

### Putative causal SNPs for GWAS regions interacting with promoters of phenotypically-relevant Mendelian disorder genes

Functional genomic datasets, such as chromatin interactions identified through Hi-C, can give us insight into the functional interpretation of GWAS variants and how they might regulate Mendelian disorder genes. Examination of chromatin interactions in human primary white adipocytes^29^ revealed further candidate credible set SNPs for metabolic traits physically interacting with promoters of phenotypically-relevant Mendelian disorder genes (**Table S9**).

Specifically, we report that a genome-wide significant SNP for BMI (rs758747, Z=6.081) physically interacts with the promoter of *CREBBP,* a gene known to cause Rubinstein-Taybi Syndrome 1 (MIM# 180849) in which obesity is one of the syndromic features^28^ (Figure 4C). These interactions can also identify the relevant isoforms of genes in disease. We identified a cluster of SNPs from the credible set of variants associated with BMI that physically interact with the promoter of a specific isoform of *CYP19A1,* a gene known to cause Aromatase Excess Syndrome (MIM# 139300) involving short stature and excess fat storage in the chest (gynecomastia)^28^ (Figure 4D). Although longer isoforms of *CYP19A1* are by default chosen to represent the gene, our data suggests that the shorter isoform is likely to be more relevant in obesity. Taken together, these results demonstrate examples of GWAS variants localizing in regulatory regions for phenotypically-relevant Mendelian disorder genes, consistent with the hypothesis that low-effect common variants contribute to complex traits by regulating genes known to cause Mendelian disorders.

## DISCUSSION

In this work we used GWAS summary statistics from 62 complex traits and genes linked to specific phenotypes within 20 Mendelian broad disorders to quantify the shared genetic basis of complex traits and Mendelian disorders. We identified a specific enrichment of phenotypically-matched and related Mendelian disorder genes in GWAS regions for complex traits; we also identified fewer pairs of complex traits and phenotypically-unmatched Mendelian disorders with similar significant enrichment. We further found that phenotypically-relevant Mendelian disorder genes are enriched for GWAS signal across complex traits, compared to all Mendelian disorder genes and other protein-coding genes. Finally, we report examples of putative causal SNPs for GWAS regions in potentially regulating phenotypically-relevant Mendelian disorder genes. We conclude with four considerations about how our results contribute to understanding of genetic architectures and biological mechanisms across complex traits and Mendelian disorders.

First, our finding of a specific enrichment of phenotypically-matched and related Mendelian disorder genes in GWAS regions for complex traits suggests that, across complex trait architectures, many complex traits share the genetic bases (and by extension, biological mechanisms) with their Mendelian forms. This supports our hypothesis that the same set of genes generally underlie both extreme and common genetic phenotypes, and suggests an important role of gene regulation by non-coding variants in complex traits. However, we note that our findings are limited by the power of each GWAS to detect significant associations. As GWAS become better-powered, we anticipate being able to identify phenotype-specific enrichments of Mendelian disorder genes in GWAS regions for more complex traits.

Second, the subset of complex trait-Mendelian disorder pairs with no known shared biology that still demonstrated significant enrichment of Mendelian disorder genes in GWAS regions can offer us novel insight into the biological mechanisms of complex traits and Mendelian disorders. A high degree of co-morbidity between complex traits and Mendelian disorders has been previously observed, regardless of phenotype-similarity^26^; these findings together suggest that many complex traits and Mendelian disorders may also be linked by the pleiotropic properties of the underlying genes, in addition to regulatory differences. These observations are also consistent with a multigenic or oligogenic architecture of human disease; the pervasive pleiotropic effects that are seen observed across complex traits are consistent with the wide-spread prevalence of multi-system, syndromic phenotypes observed across a majority of Mendelian disorders. We also confirm that LOF-intolerant genes harbor an enrichment of GWAS signal^32^; because genes with pLI > 0.9 exhibit extreme intolerance of deleterious mutation, it is possible that these genes demonstrate embryonic lethal mutant phenotypes, and are thus undiscovered as Mendelian disorder genes at this time. Our findings provide further motivation to explore phenotypic consequences of mutations in LOF-intolerant genes (particularly those enriched for GWAS signal for a particular complex trait) for phenotypically-relevant Mendelian disorders.

Third, linking Mendelian disorder genes with complex traits can help with characterization of the genetic architecture of complex traits - specifically, with genes and pathways that can be functionally characterized to identify molecular mechanisms^6^. Identifying causal variants from large-scale GWAS studies is particularly challenging given that most GWAS loci lie in non-coding regions of the genome; though thousands of genomic loci have been significantly associated with specific diseases, few casual SNPs have been functionally verified^48; 49^. Although many approaches have been used to tie a particular variant to a causal gene or genes^50–52^, including newer methods that directly link gene expression to a trait (e.g., TWAS^35^, PrediXcan^53^), we find that leveraging GWAS findings with functional data to identify candidate regulatory variants for Mendelian disorder genes can potentially lead to better interpretation of relevant genes and isoforms. Here, we demonstrate the heterogeneity of mechanisms potentially underlying causal variation, showing roles for TSS promoter regions of Mendelian disorder genes and long-range interactions involving significant GWAS regions. We expand on recent work showing that BMI-associated variants interact with genes in GWAS regions to demonstrate similar findings for Mendelian disorder genes^29^. With the appropriate functional data from relevant tissues and cell types, this phenotype-driven approach can identify relevant candidate regulatory variants and their targets. Further, from the perspective of monogenic diseases, identifying common variants that might modify the expressivity of phenotypes can provide novel insights into gene function in addition to putative drug targets. Many drugs approved by the FDA and developed by pharmaceutical companies are targeted towards the treatment of complex traits and diseases; by identifying underlying links between Mendelian disorders and complex traits through their effects on the same biological genes and pathways, we can systematically and rationally target existing drugs for complex traits and diseases towards those with rare Mendelian disorders which largely do not have any rationally targeted treatments54–56.

Last, we note that our approach of examining traits and disorders at the component-phenotype level offers us unprecedented resolution into the specific pathways involved the overall trait or disorder. In clinical medicine, genome-wide sequencing has expanded the clinical phenotypic spectrum associated with a gene ^57; 58^ through identification of pleiotropic effects due to mutations in specific protein domains^59; 60^, detected a genetic predisposition for diseases previously considered to be due to environment^13^, uncovered variable penetrance for genetic mutations previously thought to be sufficient to cause disease, and has suggested that genetic background influences the phenotypic variability of monogenic diseases^61; 62^. The phenotypic characterizations of Mendelian syndromes are deconstructed by expert clinical geneticists into component phenotypes, labeled by standardized clinical terms that identify both the primary phenotypes and phenotypes that have variable penetrance and expressivity^28; 63^. Recent work has demonstrated that incorporation of such dense phenotype information to rank putative disease-causing genetic mutations improves diagnostic rates in clinical exome sequencing tests^64; 65^. However, to our knowledge no studies as of yet have taken advantage of component Mendelian phenotypes to identify Mendelian disorders that may be phenotypically-relevant to a variety of complex traits. Ultimately, identification of GWAS-significant regions with biologically relevant genes and pathways will enable effective utilization of GWAS data in medical settings.

## SUPPLEMENTAL DATA

The supplement contains one figure and ten tables.

## ACKNOWLEDGMENTS

We would like to thank Ruth Johnson, Megan Major, Megan Roytman, Claudia Giambartolomei, Arunabha Majumdar, Jazlyn Mooney, Brendan Freund, Robert Brown, and Robert Smith for helpful discussions. This work was funded by National Institutes of Health (NIH) Training Grant in Genomic Analysis and Interpretation T32HG002536 to MKF; NIH Early Independence Award DP5OD024579 to VAA; National Institute of Mental Health of the NIH grant T32MH073526 to KB; NIH grant F31HL142180 to KMG; NIH-NCI National Cancer Institute grant T32LM012424 to DZP; and NIH grants HL-095056 and HL-28481 to PP.

## WEB RESOURCES

UCSC Table Browser: https://genome.ucsc.edu/cgi-bin/hgTables

HGNC: http://www.genenames.org/cgi-bin/download

Gene sets: https://github.com/bogdanlab/genesets

OMIM: https://omim.org/downloads/

ExAC: http://exac.broadinstitute.org/downloads

ClinVar: ftp://ftp.ncbi.nlm.nih.gov/pub/clinvar/tabdelimited/genespecificsummary.txt

